# Direct and indirect regulation of Pom1 cell size pathway by the protein phosphatase 2C Ptc1

**DOI:** 10.1101/2020.06.03.131227

**Authors:** Veneta Gerganova, Payal Bhatia, Sophie G Martin

## Abstract

The fission yeast cells *Schizosaccharomyces pombe* divide at constant cell size regulated by environmental stimuli. An important pathway of cell size control involves the membrane-associated DYRK-family kinase Pom1, which forms decreasing concentration gradients from cell poles and inhibits mitotic inducers at mid-cell. Here, we identify the phosphatase 2C Ptc1 as negative regulator of Pom1. Ptc1 localizes to cell poles in a manner dependent on polarity and cell-wall integrity factors. We show that Ptc1 directly binds Pom1 and can dephosphorylate it in vitro but modulates Pom1 localization indirectly upon growth in low glucose conditions by influencing microtubule stability. Thus, Ptc1 phosphatase plays both direct and indirect roles in the Pom1 cell size control pathway.

## Introduction

Cells coordinate growth and division to maintain cell size over generations and to adapt it to environmental conditions. The rod-shaped fission yeast *Schizosaccharomyces pombe* is widely used to dissect the molecular mechanisms of cell size control. This organism grows in length by tip extension during interphase, stops growth and divides by medial fission at a constant cell size influenced by nutrients. At the core of the decision to divide is the cyclin-dependent kinase CDK1, which drives the G2->M phase transition. CDK1 activity is directly inhibited by Wee1 kinase (Featherstone and Russell, 1991; Gould and Nurse, 1989; Lundgren et al., 1991; Russell and Nurse, 1987) and activated by Cdc25 phosphatase (Gautier et al., 1991; Kumagai and Dunphy, 1991; Russell and Nurse, 1986) and the balance between these activities controls division timing (Martin, 2009; Nurse, 1990; Wood and Nurse, 2015).

Over the past decade, it has become appreciated that Wee1 kinase integrates information from a cell geometry-sensing signaling cascade into the decision to divide. In this pathway, the DYRK-family Pom1 kinase forms plasma membrane-associated gradients with highest concentration at cell poles and serves to inhibit the SAD-family Cdr2 kinase at mid-cell (Martin and Berthelot-Grosjean, 2009; Moseley et al., 2009). Mechanistically, Pom1 directly phosphorylates Cdr2 C-terminal tail (Bhatia et al., 2014; Deng et al., 2014), which prevents Cdr2 activation by the cytosolic Ssp1 CaMKK (Deng et al., 2014). At mid-cell, Cdr2 forms stable megadalton-large membrane-associated clusters, called nodes, which also contain a second SAD-family kinase Cdr1 that phosphorylates Wee1 to inhibit its activity (Breeding et al., 1998; Kanoh and Russell, 1998; Opalko et al., 2019; Young and Fantes, 1987). Wee1 phosphorylation likely occurs during its transient visits of the nodes (Allard et al., 2018).

The spatial organization of the Pom1-Cdr2 pathway is thought to help monitor a geometric feature of cell size, coordinating cell growth with mitotic entry during the vegetative cell cycle. Recent work proposed that this feature is cell surface (Facchetti et al., 2019; Pan et al., 2014), consistent with the membrane-associated localization of all components of the pathway. As Pom1 levels at mid-cell diminish and Wee1 visits increase as cells grow, the pathway is well positioned to act as cell size sensor by linking Wee1 inhibition with cell dimensions for size homeostasis (Allard et al., 2018; Gerganova et al., 2019; Gerganova and Martin, 2018). However, as both *pom1* and *cdr2* mutants retain homeostatic capacity, this pathway may function in parallel to other homeostatic systems (Facchetti et al., 2019; Wood and Nurse, 2013). The pathway is also sensitive to changes in nutritional status. Specifically, Pom1 graded localization is altered when cells encounter low-glucose environment (Allard et al., 2019; Kelkar and Martin, 2015).

Pom1 distribution at the plasma membrane is controlled by a cycle of autophosphorylation and localized dephosphorylation (Hachet et al., 2011). Dephosphorylation of a series of autophosphorylation sites in the Pom1 cortical-binding domain by the Tea4-associated PP1 Dis2 phosphatase promotes association of Pom1 at the plasma membrane (Gerganova et al., 2019). This dephosphorylation normally takes place at cell poles, where Tea4 and its binding partner Tea1 are deposited by microtubules (Chiou et al., 2017; Martin and Arkowitz, 2014). Tea1 and Tea4 are further tethered at the membrane through direct interaction of Tea1 with the prenylated Mod5 protein (Martin et al., 2005; Snaith and Sawin, 2003), which promotes the formation of a Tea1-Mod5 polymeric network (Bicho et al., 2010). At the plasma membrane, Pom1 auto-phosphorylates to promote its return to the cytosol. It also forms cortical clusters that support its graded distribution from cell poles to mid-cell (Allard et al., 2019; Gerganova et al., 2019). These clusters, together with autophosphorylation-mediated detachment, promote robustness of the gradient (Hersch et al., 2015; Saunders et al., 2012).

Apart from the Tea4-PP1 phosphatase that controls Pom1 membrane association, no phosphatase has been shown to regulate the Pom1-Cdr2-Cdr1-Wee1 kinase cascade. To uncover potential regulatory phosphatases of the Pom1-Cdr2 network, we conducted a minimal screen for phosphatase deletions with altered cell length at division. In this work, we identify the protein phosphatase 2C (PP2C) Ptc1 as a novel interactor of Pom1 and Mod5. PP2C phosphatases are usually monomeric metal-dependent serine/threonine-specific protein phosphatases involved in the regulation of cell growth and stress signaling (Shi, 2009). In fungi, PP2C phosphatases have been strongly implicated as negative regulators of all MAPK signaling cascades (Arino et al., 2011). In fission yeast, Ptc1 negatively regulates the stress-signaling MAPK Sty1 after heat and osmotic stress through both direct dephosphorylation and indirect downstream effects (Gaits et al., 1997; Nguyen and Shiozaki, 1999; Shiozaki et al., 1994; Shiozaki and Russell, 1995). Ptc1 also acts as negative regulator of the Pmk1 cell integrity MAPK pathway (Takada et al., 2007). Ptc1 localizes to cell poles in a manner partly dependent on the SH3 domain protein Skb5, a negative regulator of the cell integrity MAPK pathway (Kanda et al., 2016; Stanger et al., 2012). Ptc1 has also recently been involved in the dephosphorylation of AMPK to regulate TOR activity in response to nitrogen limitation (Davie et al., 2015).

Here, we report that Ptc1 controls division timing through the Pom1 pathway. We show that Ptc1 localization at cell tips depends on the polarity protein Mod5 in addition to Skb5. We also report that Ptc1 moderates the previously described Pom1 gradient redistribution in low glucose conditions (Kelkar and Martin, 2015). Because Ptc1 directly binds Pom1 but exerts effects on Pom1 distribution by influencing microtubule stability and Tea4 deposition, Ptc1 likely plays both direct and indirect roles in modulating the Pom1 pathway.

## Results

### Ptc1 affects cell size at division and localizes to the cell cortex

To search for phosphatases that may regulate the Pom1-Cdr2 network, we conducted a limited screen through phosphatase deletion strains, searching for those with altered cell length at division. Because fission yeast cells grow only during interphase, cell length at division serves as a proxy for cell cycle length. Several phosphatase deletions showed altered size at division (Table S1). Here, we focus on the protein phosphatase 2C Ptc1 because its deletion and overexpression showed opposite effects on cell size at division (see below). As *ptc1* is known to be important for heat-shock response (Shiozaki et al., 1994), we first investigated whether growth of *ptc1*Δ cells was affected at different temperatures. *ptc1*Δ cells behaved similarly to wild-type cells at both high and low temperatures, as assessed by colony size formation in spot-assays (Fig. S1A). In this study, we did not investigate the effect of temperature and performed all experiments at 30°C, unless stated otherwise, to probe the role of Ptc1 in cell size regulation.

### Ptc1 localization to the cell cortex depends on Skb5 and Mod5

Previous work suggested that Ptc1 localizes to the cell poles as well as in the cytoplasm and cell nucleus (Kanda et al., 2016). We confirmed these observations by tagging of *ptc1* at the endogenous locus with GFP, which yielded a largely functional protein, as assessed by cell length measurements (Table S1; also see Fig 3B below). Ptc1-GFP was indeed observed at the cell cortex and at the septum of dividing cells, as well as in the cytoplasm and weakly in the nucleus (Fig. 1A, Movie 1). Quantitative measurement of Ptc1-GFP cortical signal revealed higher intensities at the cell tips than at cell middle (Fig. 1A, right panel), indicating a shallow gradient-like pattern.

**Figure 1:**
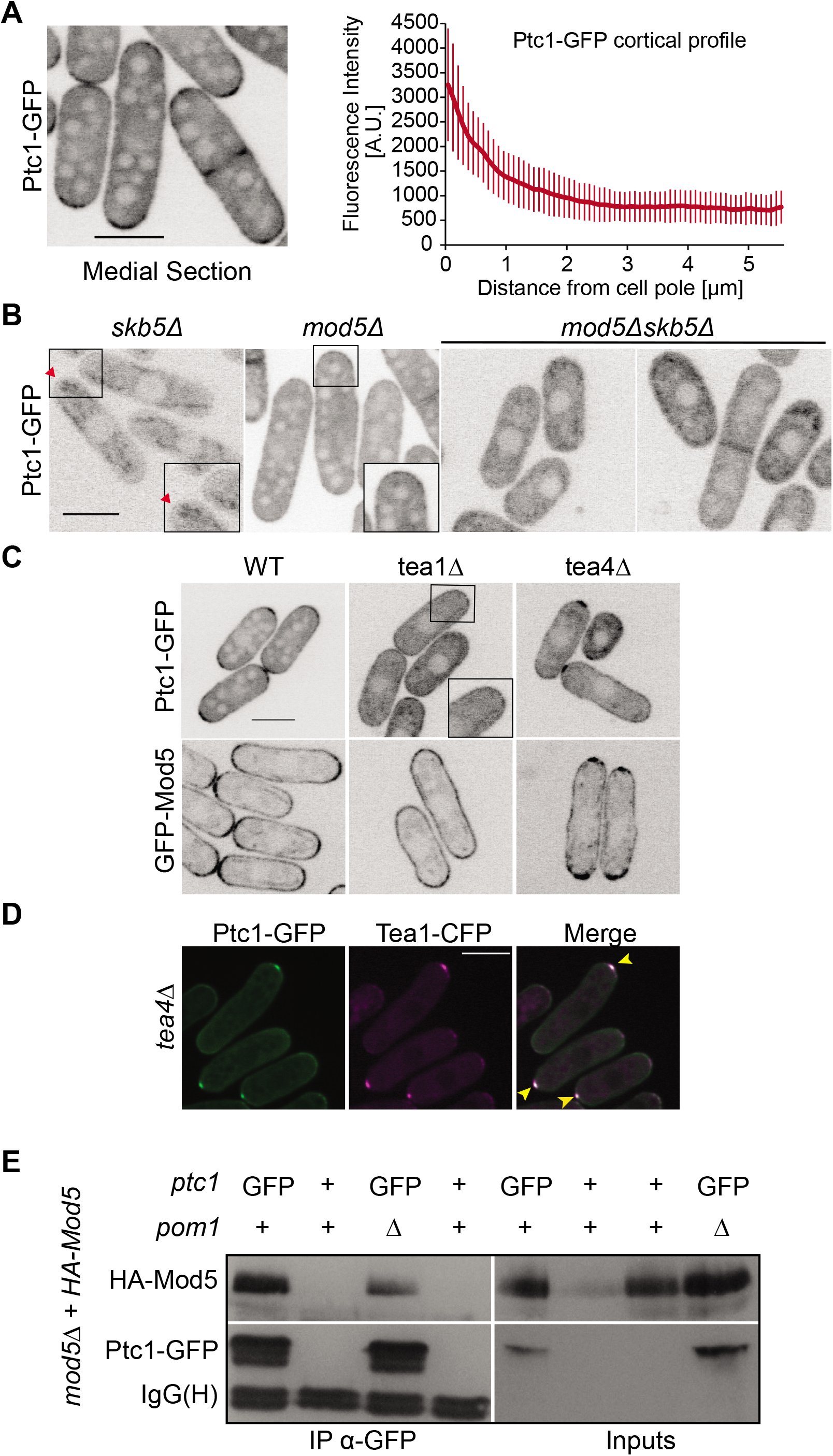
Ptc1 localizes to the cell cortex. **A.** Localization of Ptc1-GFP in fission yeast cells. Left: A medial section of confocal microscope images. Right: Average cortical profile of Ptc1-GFP (n=20 cells). 5 images were acquired over time at maximum speed and fluorescence intensities were measured on sum projections from cell tip to cell tip using 3-pixel wide segmented line scan in ImageJ. Error bars are standard deviations. **B.** Ptc1-GFP localization in *skb5Δ, mod5Δ* and *mod5Δskb5Δ* mutants. Insets show magnifications of selected cell poles. **C.** Ptc1-GFP and GFP-Mod5 localization in *tea1Δ* and *tea4Δ*. **D.** Co-localization of Tea1-CFP and Ptc1-GFP (acquired in the YFP channel) in *tea4*Δ cells. Yellow arrowheads indicate colocalizing Tea1 and Ptc1 dots. False color image of Tea1-CFP. **E.** Co-immunoprecipitation of HA-Mod5 and Ptc1-GFP from *mod5*Δ cells expressing HA-Mod5 on a plasmid under the inducible *Pnmt1* promoter. Ptc1-GFP was immunoprecipitated using anti-GFP antibody and co-immunoprecipitation of Mod5 was revealed using anti-HA antibody. IgG(H) serves as the immunoprecipitation control. Inputs are shown on the right. Note that the last two lanes in the input blot are inverted compared to the IP blot. Images are sum projections of 5 confocal images acquired over time. All scale bars: 5μm.

To uncover factors responsible for Ptc1 localization, we first investigated the possible role of Skb5, previously shown to bind Ptc1 (Kanda et al., 2016; Stanger et al., 2012). Skb5 is an SH3 domain protein, named after its <u>S</u>hk1 <u>k</u>inase <u>b</u>inding function (Yang et al., 1999), which was shown to regulate cell integrity Pmk1 MAPK signaling by controlling the localization of the MAPKKK Mkh1 (Kanda et al., 2016). Skb5-GFP itself weakly localizes to cell tips (Fig. S2A) (Kanda et al., 2016) when expressed from its endogenous locus. In *skb5Δ* cells Ptc1 became largely cytosolic, though faint amounts of Ptc1 could still be detected at cell tips (Fig. 1B). Thus, Skb5 is one important contributor to Ptc1 localization to cell tips.

We investigated other possible determinants of Ptc1 cortical localization. Ptc1 localization was not affected by acute depolymerization of the actin or microtubule cytoskeleton (Fig. S2B). By contrast, the loss of polarity landmarks Tea1 (Mata and Nurse, 1997), Tea4 (Martin et al., 2005; Tatebe et al., 2005), or Mod5 (Snaith and Sawin, 2003) altered Ptc1 localization (Fig 1B-C). Deletion of Tea3 (Arellano et al., 2002) or Pom1 had minor effects (Fig S2C). Deletion of Bud6 (Glynn et al., 2001) produced no detectable change in Ptc1 distribution (Fig S2C). Several observations pointed to Mod5 acting as a direct Ptc1 link to the membrane. First, deletion of *mod5* had the strongest effect on Ptc1 localization (Fig 1B): in *mod5*Δ Ptc1-GFP localization ranged from completely cytosolic to a faint uniform cortical signal in some cells. Second, Ptc1-GFP was spread more evenly around the cell cortex in *tea1*Δ cells, in which Mod5 is also not restricted to cell poles (Fig 1C) (Snaith and Sawin, 2003). Third, Ptc1-GFP formed a dot-like structure at the non-growing end of *tea4Δ* cells, where Tea1 and Mod5 accumulate (Fig 1C) (Martin et al., 2005). Indeed, in *tea4*Δ cells Ptc1-GFP co-localized with Tea1 (Fig 1D). Furthermore, the Ptc1-GFP dot was completely lost upon additional deletion of *mod5* (Fig S2C). Conversely, *ptc1*Δ did not affect the localization of Tea1, Tea4 or Mod5 (Fig S2D). Thus, the prenylated protein Mod5 may serve as an anchor for Ptc1 at the cell cortex, similar to its role in anchoring Tea1 (Snaith and Sawin, 2003).

To test whether Mod5 associates with Ptc1, we performed co-immunoprecipitation experiments, using cells expressing HA-Mod5 from an inducible plasmid and Ptc1-GFP from the endogenous locus. Mod5 co-immunoprecipitated with Ptc1 from cell extracts (Fig 1E). We also determined that the Mod5-Ptc1 association is independent of Pom1 (Fig 1E and S2E; see below). We conclude that Mod5 forms a complex with Ptc1 and can dictate its binding at the cell cortex additionally to Skb5.

Finally, since single Skb5 and Mod5 are two Ptc1 interactors that control its localization, we probed the relationship between these two landmarks. In *skb5Δ mod5Δ* double mutant cells, Ptc1 was fully cytosolic in interphase cells (Fig 1B), suggesting that each landmark contributes to the residual cortical localization observed in single mutant cells. We note that in dividing cells Ptc1 retained the ability to localize at the septum, indicating the existence of at least a third landmark (Fig 1B). Mod5 and Skb5 do not play a role in each other’s localization, as no major changes of distribution of GFP-Mod5 were observed in *skb5Δ* and vice versa (Fig S2A). These data establish that Ptc1 localizes to cell poles through interaction with two independent landmarks, Skb5 and Mod5.

### Ptc1 regulates cell size through Pom1 kinase

The slightly longer size at division of *ptc1*Δ cells and the cortical localization of Ptc1 prompted us to further probe its role in cell size regulation in relation with the cortical kinase Pom1. In exponentially growing *ptc1*Δ cells, calcofluor-staining revealed a slight yet significant increase of cell length at division (Fig 2A). Other phenotypes such as septum positioning and polarized growth were not significantly different from the wildtype (Fig S1B). Conversely, overexpression of *ptc1* from the inducible *p^nmt1^* promoter integrated at the endogenous locus led to a significant reduction in cell length at division (Fig 2A), consistent with previous data (Gaits et al., 1997), suggesting that *ptc1* influences the timing of entry into mitosis in fission yeast cells. Interestingly, *ptc1* deletion and overexpression had no effect on cell size at division in *pom1*Δ cells (Fig 2A). Therefore, *pom1*Δ is epistatic to *ptc1*Δ, suggesting that Ptc1 regulates the timing of mitotic entry through Pom1. These data establish the phosphatase Ptc1 as a novel regulator of the Pom1 pathway in fission yeast cells.

**Figure 2:**
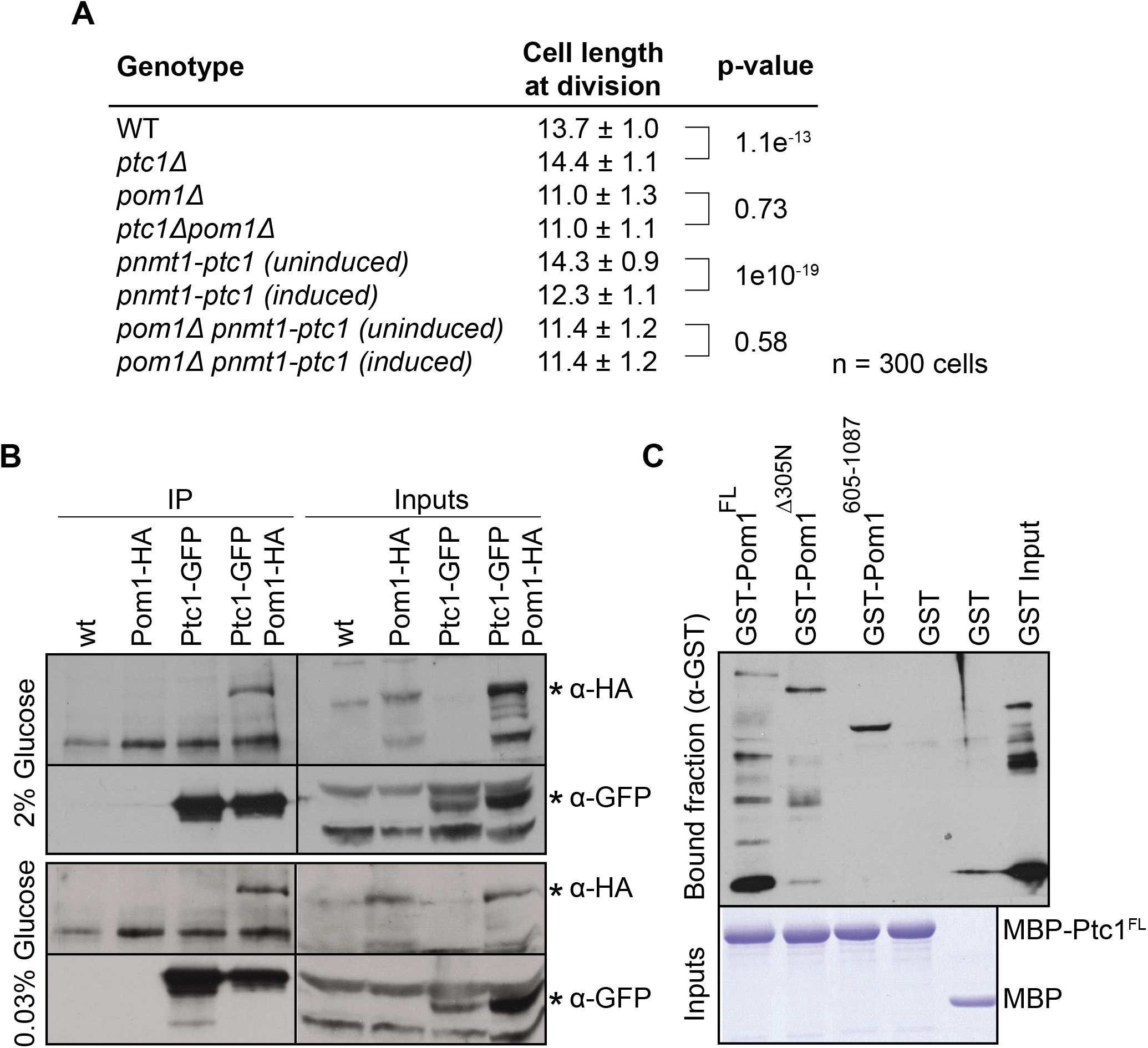
Ptc1 localization to the plasma membrane is dependent on Skb5 and Mod5. **A.** Cell lengths at division quantified from >300 cells in at least 3 independent experiments, statistical significance measured by student t-test against either wild type or uninduced counterpart. **B.** Co-immunoprecipitation of Ptc1-GFP and Pom1-HA. Protein extracts were made from cells grown in 2% glucose for 24h and from cells shifted to 0.03% glucose for 2h. 2mg whole cell extract was used for immunoprecipitation assay. 100μg inputs were loaded. **C.** Pom1 binds Ptc1 *in vitro*. Bacterially expressed proteins assayed by affinity columns. GST tagged Pom1^FL^ and fragments Pom1^Δaa305N^ and Pom1^aa605-1087^ tested for interaction with recombinant MBP tagged Ptc1 ^FL^.

Consistent with the idea that Ptc1 regulates Pom1, we found that the two proteins form a complex both *in vivo* and *in vitro. In vivo*, Pom1-HA co-immunoprecipitated with Ptc1-GFP in several growth conditions (Fig 2B). *In vitro*, we found that recombinant GST-Pom1 directly binds recombinant MBP-Ptc1 (Fig 2C). This interaction was independent of Pom1 N-terminal membrane-binding region. Thus, Ptc1 phosphatase and Pom1 kinase bind directly to each other and Ptc1 modulates cell length at division in a manner dependent on Pom1.

### Ptc1 activity is required for function and localization

We investigated the role of Ptc1’s phosphatase activity. To verify if Ptc1 works as an active phosphatase *in vitro*, we purified GST-Ptc1 from bacterial cells and used it in an *in vitro* phosphatase assay with recombinant, autophoshorylated Pom1 as substrate. Recombinant Pom1 runs as a smear because of autophosphorylation and shows a clear shift in mobility upon dephosphorylation by the phosphatase PP1 (Hachet et al., 2011). Ptc1 induced a similar shift in Pom1 mobility (Fig. 3A), suggesting that Ptc1 is active as a phosphatase *in vitro*. Through sequence alignment with Ptc1 homolog in *Saccharomyces cerevisiae*, we first identified 4 amino acids (aa; D109, G110, D275, and D314) within the phosphatase domain of fission yeast Ptc1 predicted to be involved in divalent metal ion binding and therefore important for catalytic activity. Mutation of one or several aa identified above, yielding Ptc1^1A^ (D275A) and Ptc1^3A^ (D109A/G110A/D314A) prevented the shift in mobility (Fig 3A, Fig S3) confirming the phosphatase activity of Ptc1. We note that Pom1 activity towards its substrate Cdc15 (Bhattacharjee et al., 2020) was not dramatically affected by dephosphorylation, though we cannot exclude subtle changes. The Ptc1^3A^ catalytic-dead mutant was used further in the study.

**Figure 3:**
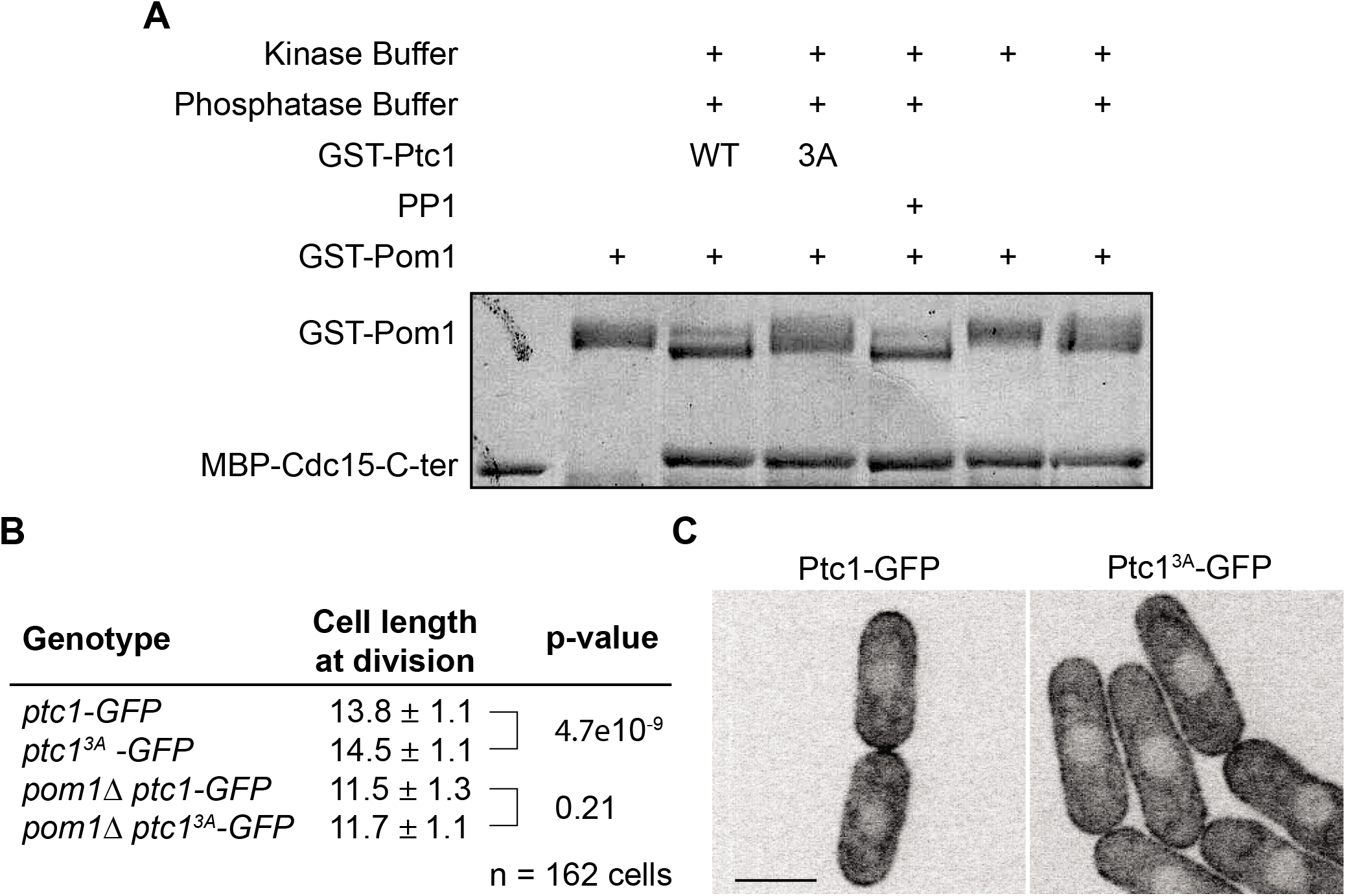
Ptc1 dephosphorylates Pom1 *in vitro* and its activity contributes to its localization. **A.** In vitro phosphatase assay, using recombinant proteins. Autophosphorylated recombinant GST-Pom1 was used as substrate and incubated with recombinant GST-Ptc1 (WT or 3A mutant) or commercial PP1 as indicated. Autophosphorylated Pom1 migrates as a smear, which collapses to two fast-migrating bands upon dephosphorylation. Phosphatase-treated Pom1 was then used in a kinase assay to monitor Pom1 activity. Recombinant MBP-Cdc15C was used as Pom1 substrate, as this fragment is quantitatively phosphorylated by Pom1 leading to slower migration (Bhattacharjee et al., 2020). Note that the phosphorylation status of Pom1 does not alter its activity towards Cdc15C. **B.** Cell lengths at division quantified from >160 cells in 3 independent experiments, statistical significance measured by student t-test against wild type or *pom1Δ*. **C.** Localization of Ptc1-GFP and Ptc1^3A^-GFP. Scale bar: 5μm.

To test the role of phosphatase activity *in vivo*, we introduced a *ptc1^3A^-GFP* allele as a sole copy of *ptc1* at the endogenous *ptc1* locus. *ptc1^3A^-GFP* cells phenocopied *ptc1*Δ mutants: these cells were slightly, but significantly, longer at division than wildtype cells, and *pom1*Δ was epistatic over this phenotype (Fig 3B). Interestingly, Ptc1^3A^-GFP failed to be enriched at cell poles and instead spread all along the cortex (Fig 3C). Therefore, the catalytic activity of Ptc1 is critical for its function in cell size control as well as for its cell tip enrichment.

### Localization and function of Ptc1 in glucose-limited conditions

Ptc1 is involved in the regulation of MAPK pathway in fission yeast under different kinds of environmental stresses (Gaits et al., 1997; Nguyen and Shiozaki, 1999; Shiozaki et al., 1994; Shiozaki and Russell, 1995; Takada et al., 2007). To further understand its role in cell size regulation, we tested if Ptc1 itself is affected under stress, and if it is involved in cell size regulation under these conditions. We first tested the localization of Ptc1 in cells subjected to osmostress (sorbitol and KCl), nutritional stress (lack of nitrogen and low glucose conditions), and high temperature. Under all conditions except low glucose, Ptc1 remained well localized at the cortex (Fig S4A-B). In low glucose (0.08% glucose) Ptc1 was evenly spread around the cell cortex (Fig 4A). Ptc1-GFP intensity measurements around the cell cortex indeed revealed a clear spread of signal from cell tips to the cell middle in 0.08% glucose growth conditions.

**Figure 4:**
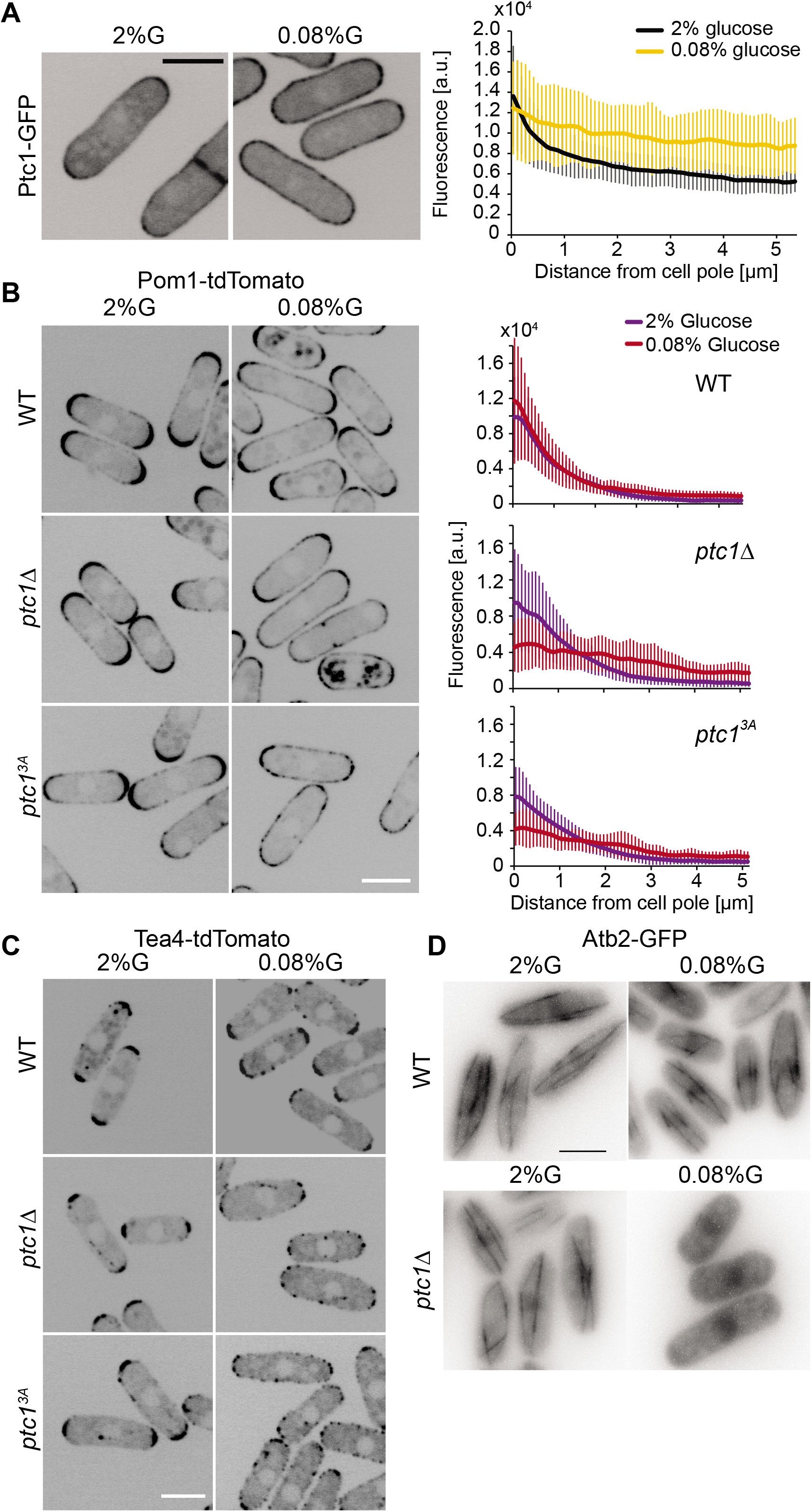
Ptc1 keeps Pom1 from spreading to the cell sides in limited glucose conditions. **A.** Localization of Ptc1-GFP in EMM-ALU 2% glucose and 0.08% glucose. Graph on right shows average profiles of wild type Ptc1-GFP from n=20 cells. Individual profiles are shown in Fig S4B. Error bars are standard deviations. **B.** Localization of Pom1-tdTomato in wt, *ptc1*Δ and *ptc1^3A^-GFP* (GFP not shown). Graphs on right show average fluorescence profiles of Pom1-tdTomato obtained from individual cells (N=20). Error bars are standard deviations. **C.** Localization of Tea4-tdTomato in wt, *ptc1*Δ and *ptc1^3A^-GFP* backgrounds (GFP not shown). **D.** Atb2-GFP signal in wild type and *ptc1*Δ backgrounds. Widefield microscopy. The indicated time is the exact time point during the imaging interval. In all panels, cells were grown in 2% glucose (G) for 24h and shifted to 0.08% glucose for 1h before imaging. In (A-C), images are sum projections of 5 confocal images acquired over time. In (D), the images shown were taken 1h30 (top) and 1h10 (bottom) after transfer to 0.08% glucose. Scale bars: 5μm.

Because of the epistasis of *pom1*Δ over *ptc1*Δ and because Pom1 localization is itself altered in low glucose conditions, we tested if loss or inactivation of Ptc1 affected Pom1 localization. Indeed, our previous work had shown that complete glucose starvation (≤0.03%) causes a uniform redistribution of Pom1 around the cell cortex, whereas limiting glucose conditions (0.08%) cause a much milder redistribution of Pom1 to the cell middle (Kelkar and Martin, 2015). While loss of Ptc1 or its activity had no major effect on Pom1 localization in 2% glucose (Fig 4B), it had a significant effect in 0.08% glucose: in *ptc1*Δ and *ptc1^3A^* cells, Pom1 nearly completely lost its gradient shape and displayed in an almost flat profile around the cortex as compared to wild type (Fig 4B, right top panel). Thus, loss of Ptc1 function exacerbates the redistribution of Pom1 in low glucose conditions.

Pom1 localization to the cell cortex depends on local dephosphorylation by the Tea4-Dis2 PP1 complex. In glucose-rich conditions, Tea4 is deposited by microtubules at cell poles, where it nucleates the formation of Pom1 concentration gradients (Hachet et al., 2011). In glucose-poor conditions, destabilization of MTs leads to Tea4 delivery at the cell sides, resulting in Pom1 re-localization around the cell periphery (Kelkar and Martin, 2015). We thus asked whether Ptc1 controls Tea4 deposition and/or microtubule organization. In *ptc1*Δ and *ptc1^3A^-GFP*, Tea4 localized correctly at cell poles in 2% glucose. By contrast, in glucose-limited conditions (0.08%), Tea4 was strongly delocalized from the poles with Tea4 dots spread around the cortex (Fig 4C). We made similar observations regarding microtubules: In 2% glucose no major differences in microtubule dynamics between the wild type and the mutant were observed, but exposure of *ptc1Δ* cells to 0.08% glucose conditions destabilized microtubules. Indeed we found that, 30min to 1h after transfer to 0.08% glucose, microtubules were depolymerized in *ptc1*Δ cells, but not in wildtype cells (Fig 4D). In summary, these data suggest that destabilization of microtubules in *ptc1Δ* cells leads to aberrant Tea4 deposition at cell sides and subsequent loss of the Pom1 gradient shape.

## Discussion

In this work, we present an initial dissection of the mode of localization and action of the PP2C phosphatase Ptc1 in regulating cell size. In agreement with previous data, we have found that Ptc1 promotes division at a shorter cell size, as shown by the extended length at division of cells carrying *ptc1*-deletion and -inactive alleles and the shorter length of cells overexpressing it. Our genetic data indicate that it does so through Pom1. What may be the mechanism?

A first simple scenario would be that Ptc1 serves to reverse Pom1 phosphorylation. This possibility is evoked from our discovery that Ptc1 forms complexes with Pom1 *in vivo* and directly binds and dephosphorylates it *in vitro*. In previous work, we showed that Pom1 gradients are patterned by a cycle of phosphorylation and dephosphorylation, where dephosphorylation of Pom1 by the Tea4-associated PP1 phosphatase within Pom1’s membrane-binding region promotes membrane attachment, and autophosphorylation reduces membrane binding (Gerganova et al., 2019; Hachet et al., 2011). If Ptc1 was acting to dephosphorylate these sites, one would expect that in *ptc1*Δ cells Pom1 exhibits higher phosphorylation and reduced membrane-binding, as in *tea4*Δ cells. However, our observations are not consistent with this prediction: in glucose-rich conditions, Pom1 distribution was not significantly altered, and in glucose-limiting conditions Pom1 showed an opposite phenotype, decorating the entire cell cortex. We conclude that Ptc1 is highly unlikely to dephosphorylate Pom1 within the cortex-binding region. As Pom1 autophosphorylates on nearly 40 other serines and threonines (Hachet et al., 2011), one possibility is that Ptc1 reverses other phosphorylation events. The opposite cell size phenotypes of *ptc1*Δ and *pom1*Δ predicts that dephosphorylation by Ptc1 may serve to reduce Pom1 activity.

A second scenario is that Ptc1 controls Pom1 indirectly. Although Ptc1 likely does not modulate Pom1 localization through dephosphorylation of the Pom1 cortex-binding domain, Ptc1 clearly influences Pom1 distribution. Indeed, in glucose-limiting conditions, where Pom1 gradient distribution is less robust due to destabilization of microtubules, deletion of *ptc1* caused a strong re-distribution of Pom1 around the entire cell cortex. We hypothesize that this re-localization reflects an indirect function of Ptc1 on Pom1 distribution. Our previous work had shown that microtubule destabilization in glucose-limited conditions leads to deposition of Tea4 and consequent Pom1 re-distribution at cell sides (Kelkar and Martin, 2015). The microtubule destabilization relies on PKA inhibiting CLASP-dependent microtubule rescue. We have now observed that *ptc1*Δ exacerbates the microtubule destabilization phenotype, leading to Pom1 re-distribution. While additional work will be required to understand how Ptc1 exerts this function, it may for instance counteract PKA-dependent phosphorylation.

The observation that Ptc1 itself localizes at the plasma membrane suggests this may be its place of action to modulate the Pom1-Cdr2 geometry-sensing cell size pathway. Ptc1 is somewhat enriched at cell poles dependent on several factors. We found that Ptc1 recruitment to the cell cortex requires two proteins that Ptc1 associates with: the prenylated protein Mod5 and the SH3-domain protein Skb5 (this work and (Kanda et al., 2016)). Ptc1 may thus integrate information from the polarity/microtubule machinery through Mod5 and from the cell wall/growth status through the cell integrity pathway dependent Skb5 protein. Ptc1’s restriction to cell poles also depends on its own activity. These observations raise the question of the importance of Ptc1 localization for its cell size-modulatory function. We can speculate that the increased amounts of Ptc1 at cell poles may skew stabilization of microtubules towards cell poles, thus favoring the delivery of Tea4 at these locations. Similarly, they may help delay Pom1 autophosphorylation that promotes its detachment by lowering Pom1 activity at these locations. The identification of Ptc1 as a first phosphatase regulating Pom1 adds to a growing number of regulatory, cortical factors modulating the Pom1-Cdr2 geometry-sensing cell size pathway, such as for instance the methyl transferase Skb1 (Deng and Moseley, 2013). These regulators, themselves important modulators of stress-signaling cascades, may help cells adapt their size to environmental stimuli.

## Acknowledgements

This work was supported by Swiss National Science Foundation research grants (31003A_138177, 310030B_176396 and CRSII3_160728) to SGM.

## Materials and Methods

### Yeast genetics and culture

*Schizosaccharomyces pombe* strains obtained from genetic crosses were selected by tetrad dissection and replica plating with appropriate supplements or drugs. All transformations were performed using the lithium acetate-DMSO method. Tagged and deletion strains were constructed by using a PCR-based approach (Bähler et al., 1998) and confirmed by PCR. Ptc1 overexpressing strains were obtained by introducing *Pnmt1* promoter upstream of *ptc1* at its endogenous locus using PCR-based approach. All strains were genotyped using PCR.

Cells were grown in standard Edinburgh minimal media (EMM) with appropriate supplements adenine, leucine and uracil when required. For glucose limitation assays, cells were grown in 2% glucose to mid-log phase, washed three times with EMM 0.08% glucose and incubated in the same medium before imaging. For cell size experiments, cells were grown in either EMM 2 % glucose or EMM 0.08% glucose for 24h before imaging.

For expression of HA-Mod5, yeast cells were transformed with pREP41-HA-Mod5 plasmid and selected on EMM supplemented with adenine and uracil. Cells were grown in EMM adenine, uracil, and with or without thiamine (5 μg/ml) at 30°C for 24h before pelleting down for western-blot or immunoprecipitation experiments.

For overexpression from *nmt1* promoters, cells were grown for 24h at 30°C in EMM supplemented with adenine, uracil, leucine with or without thiamine (15μM).

For cell size measurement, Calcofluor (Sigma) was added at a final concentration of 5 μg/ml from a 200X stock solution.

### Plasmid construction

For GST-Ptc1, Ptc1 amplified from gDNA was cloned into pGEX-4T-1 GST vector between EcoR1 and Xho1 sites. Mutant GST-Ptc1 plasmids were generated by site-directed mutagenesis on GST-Ptc1 plasmid. Mod5 was cloned in a pREP41-HA (N) vector between Sal1 and BamH1 sites for the production of HA-Mod5. All plasmids constructed were confirmed by sequencing.

### Microscopy

Images on live cells were acquired at room temperature, except stated otherwise, either on a Spinning Disk confocal microscope or on DeltaVision epifluorescence microscope. Confocal images were acquired using a Perkin Elmer Leica DMI4000B inverted microscope equipped with an HCX PL APO ×100/1.46 numerical aperture (NA) oil objective and a PerkinElmer Volocity Confocal system spinning disk microscope including a Yokagawa CSU22 real-time confocal scanning head, an solid-state laser lines and a cooled 14-bit frame transfer EMCCD C9100-50 camera, as described (Bendezu et al, PLoSONE 2012). Stacks of z-series confocal sections were acquired at 0.3 μm intervals with the Volocity software. For the rest of the images, 5 images were acquired over time. Widefield microscopy was performed on conventional imaging mode of a DeltaVision OMX SR imaging system, equipped with a 60x 1.49 NA TIRF oil objective (oil 1.514) a front illuminated sCMOS camera size 2560 x 2160 pxl (manufacturer PCO).

### Image analysis

Fluorescence intensity quantifications shown in Fig.4 and S4 were performed on individual cells on a sum projection of spinning disk confocal images (5 images acquired over time). For measurement of fluorescence intensity along the cell cortex, 3-pixel wide segmented line in ImageJ was used to draw a line by hand at the periphery of the cell and fluorescence intensity obtained using the Analyze<Measure tool. The fluorescence intensity measured was corrected for background fluorescence intensity (measured just outside the cell examined).

Cell length measurements were made on DeltaVision acquired calcofluor stained images. Line segment tool in ImageJ was used to measure the distance between cell ends of dividing cells.

### Spot Assays

Cultures were adjusted to the same OD600 for the highest concentration sample and subsequent 10-fold serial dilutions were made in a total of 200μL. 5 μL of diluted culture was spotted on YE plates are grown at respective temperatures.

### Recombinant protein production

Expression of GST-Pom1, GST-Ptc1 and GST alone was induced in BL21 bacteria from the pGEX-4T-1-derived plasmids as described above. Expression of MBP-Ptc1 induced from a pMAL derived vector. Briefly, cells were grown overnight in 25ml LB with 100μg/ml ampicillin at 37°C. 250ml of LB-ampicillin was inoculated with 6.25ml of the saturated culture, grown 3 h at 37°C. Protein expression was induced by the addition of 100μM IPTG for 5 h at 18°C. Bacterial pellets were resuspended in 10ml cold PBS and sonicated 3 times for 30s each (50% amplitude). The sonicates were incubated with 1% TritonX-100 at 4°C, and centrifuged 15min at 4°C at 10,000g. Soluble extract was incubated with 200 μl of glutathione sepharose beads (50% slurry) for 2h at 4°C. Finally, beads were washed 3 times with cold PBS, and eluted in three steps in 100 μl elution buffer (15mM reduced glutathione in 50 mM Tris-HCl, pH 8) elution buffer.

### Immunoprecipitation and Western Blotting

For western blots and immunoprecipitation assays, whole cell extracts (WCE) were prepared in CXS buffer (50 mM HEPES, pH 7.0, 20 mM KCl, 1 mM MgCl, 2 mM EDTA, pH 7.5 and protease inhibitor cocktail, Roche) from cells grown in EMM with appropriate glucose concentrations and with or without thiamine (5μg/ml), using glass beads and a bead beater. Total protein concentration was measured using standard Bradford assay. 2mg WCE was used for immunoprecipitation. Briefly, 50μl Dynabeads (Thermofisher) per sample was washed 3 times with cold 1X PBS and 2μg anti-GFP monoclonal antibody (Roche) per sample was conjugated with Dynabeads in 1ml cold PBS at 4°C for 4h. Antibody-conjugated beads were then washed 2 times with cold PBS and once with CXS buffer. In low-protein binding tubes, 2mg WCE was combined with 50μl antibody-conjugated beads in 1ml CXS buffer supplemented with 100mM NaCl and 0.1% NP-40 and left at 4°C for overnight on a rotation wheel. Next day, any unbound antibody was removed by washing the WCE 3 times with CXS with 0.1% NP-40 and increasing amounts of NaCl (100mM, 150mM and 200mM, respectively). Samples were boiled with 1X Laemmli buffer and 100μg of WCE were loaded together with the immunoprecipitates on SDS-PAGE gels. Bands were revealed using anti-GFP polyclonal antibody (SantaCruz) and mouse anti-HA monoclonal antibody (HA.11; Covance).

### *In vitro* phosphatase assay

*In vitro* phosphatase assay was performed using purified GST-Pom1 and GST-Ptc1 wild-type and GST-Ptc1 mutant proteins. Briefly, 200ng GST-Pom1 was incubated with 1μg GST-Ptc1, GST-Ptc1 mutants, or commercial PP1 (NEB #P0754S) in phosphatase buffer with 5X composition: 250mM Tris-Cl pH7.5, 1mM EDTA, 0,5% β-mercaptoethanol, 100mM MgCl_2_ at 37°C for 1.5h. The reaction was stopped using 1X Laemmli buffer. Samples were boiled at 95°C for 5 mins before loading on SDS-PAGE gel and stained with silver stain. The gel was run slowly at 80V for 3h to enable Pom1 shift. Simultaneously, 1μg GST-Ptc1 and GST-Ptc1 mutants were loaded on another SDS-PAGE gel for Coomassie staining.

---

**Figure S1:**
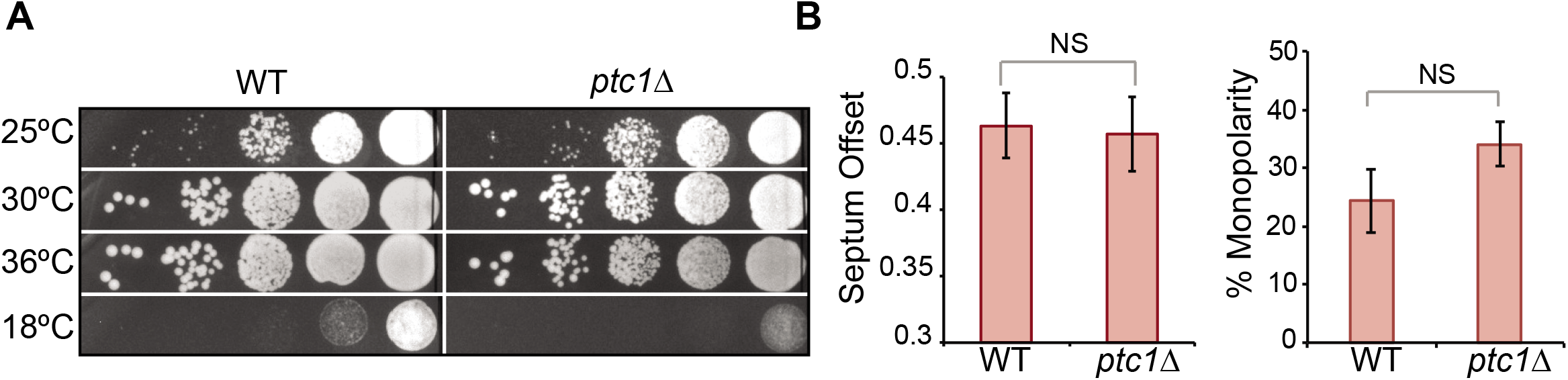
*ptc1*Δ cells are not temperature sensitive. **A.** Serial dilution assay of wild type and *ptc1*Δ cells at different temperatures. **B.** Septum offset quantification in wild type and *ptc1*Δ cells. More than 120 cells per strain were counted from 2 independent experiments. Right: percentage of monopolar cells from wild type and *ptc1*Δ strains. More than 100 septated cells were counted from 3 independent experiments. *p-values* determined using student t-test.

**Figure S2:**
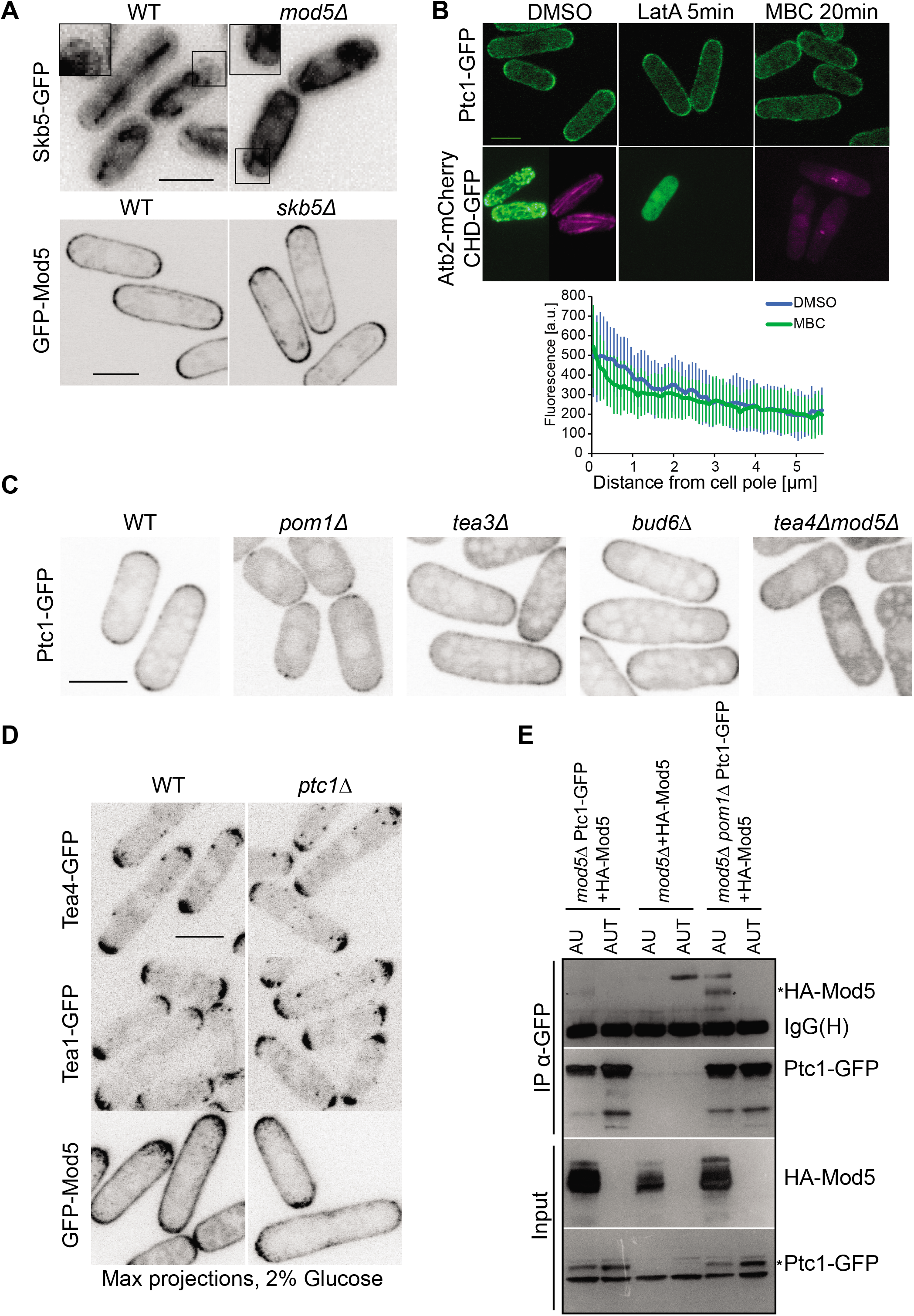
Investigation of Ptc1 localization. **A.** Localization of Skb5-GFP in WT and *mod5Δ* cells and GFP-Mod5 in WT and *skb5Δ* cells (widefield microscopy). Insets show magnifications of selected cell poles. **B.** Localization of Ptc1-GFP in MBC and LatA treated cells. Top: Cells were treated with DMSO, 200μM LatA for 5’, and 25μg/ml MBC for 20’ and imaged using confocal microscope. Max projections of 14 z-slices are shown. Bottom: Cells with labeled mCherry-Atb2 and CHD-GFP were treated and imaged similarly as above to serve as controls for effective depolymerization of MTs and F-actin upon drug treatment. LatA treatment effectively removed actin cables and patches. MBC treatment effectively destroyed interphase MTs. Bottom: average profile from quantification of Ptc1-GFP fluorescence intensity along the cortex of 16 cells treated with MBC or DMSO control for 20min. Quantifications done on sum projections of 14 z-slices. Error bars are standard deviations. **C.** Localization of Ptc1-GFP in WT, *pom1Δ, bud6Δ, tea3Δ*, and *tea4Δmod5Δ* cells. **D.** Localization of Tea4-GFP, Tea1-GFP and GFP-Mod5 in WT and *ptc1Δ*. **E.** Co-immunoprecipitation of HA-Mod5 and Ptc1-GFP from *mod5*Δ and *mod5Δ pom1*Δ cells expressing HA-Mod5 on a plasmid under the inducible *Pnmt1* promoter. Ptc1-GFP was immunoprecipitated using anti-GFP antibody and co-immunoprecipitation of Mod5 was revealed using anti-HA antibody. IgG(H) serves as the immunoprecipitation control. Inputs are shown on the bottom. Note there are a couple of background bands migrating more slowly than HA-Mod5. Scale bars: 5μm.

**Figure S3:**
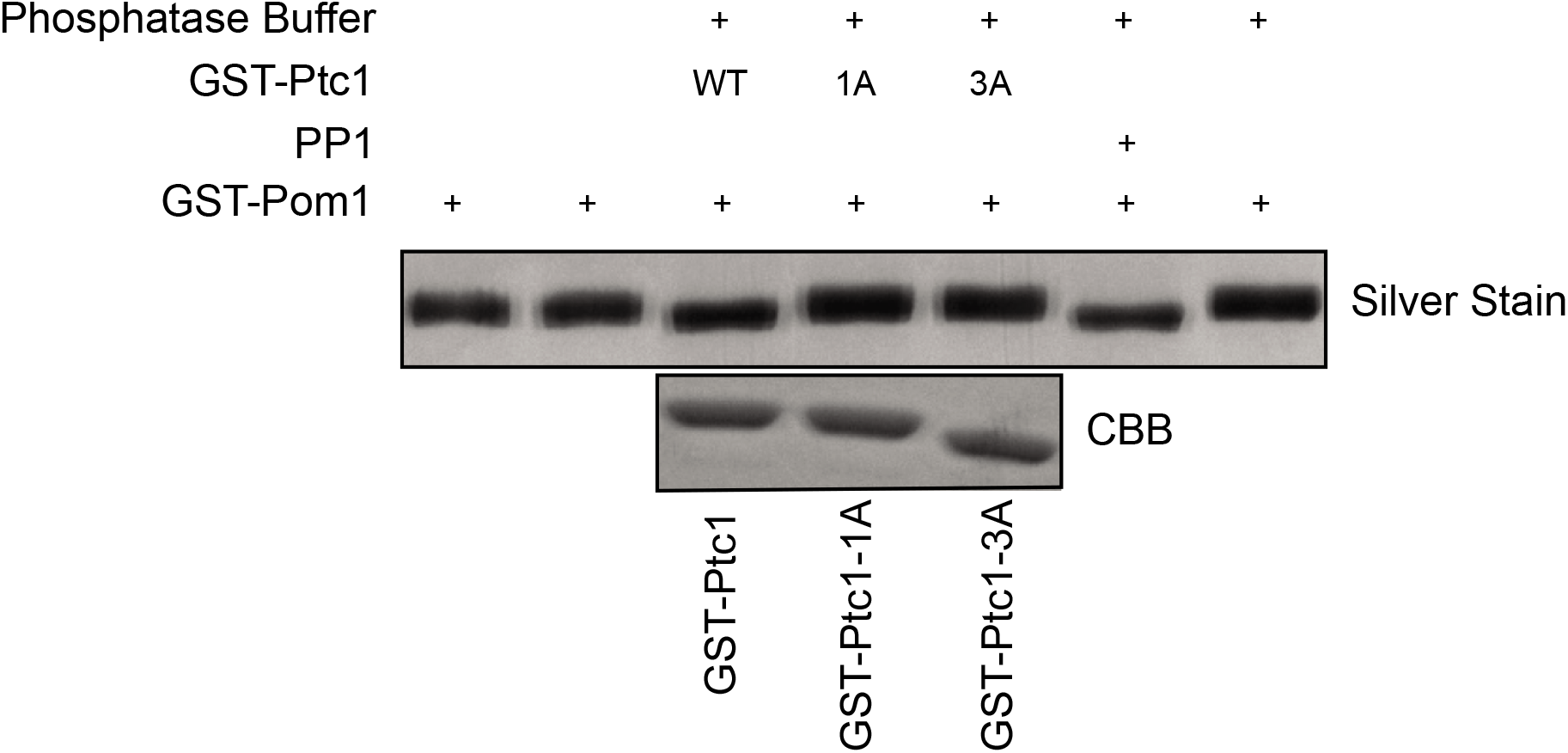
Ptc1 phosphatase activity in vitro. In-vitro phosphatase assay. Purified proteins were incubated in phosphatase buffer at 37°C for 1.5h. Top: GST-Pom1 shift revealed using silver staining. Bottom: Loading control for GST-Ptc1 proteins revealed using Coomassie blue staining (CBB).

**Figure S4:**
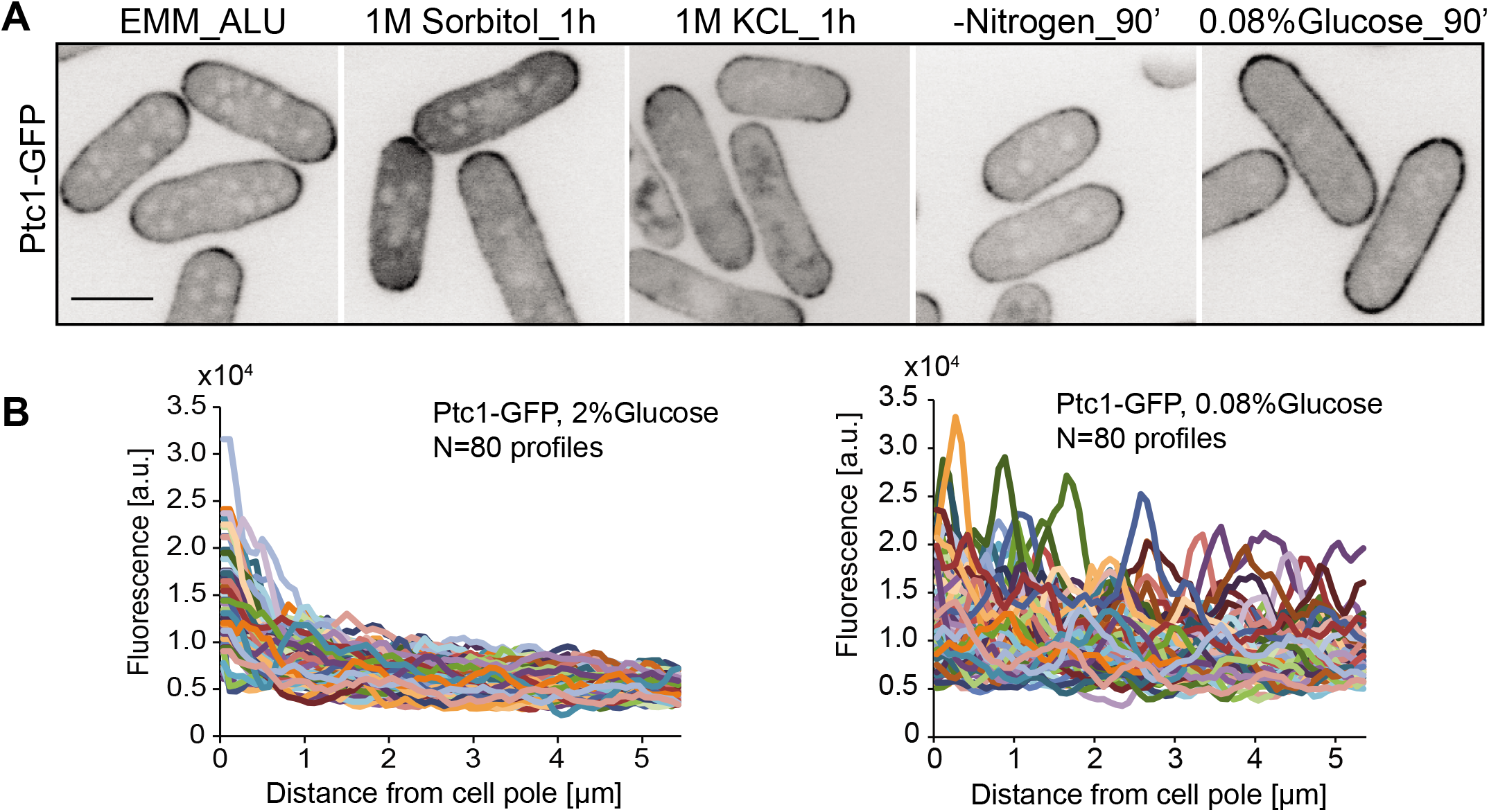
Controls for Pom1 and Ptc1 localization. **A.** Localization of Ptc1-GFP in different stresses. Cells were grown in EMM-ALU for 24h and then treated with 1M sorbitol, or 1M KCL for 1h before imaging. For nutrient starvation, cells grown in EMM-ALU were washed 3 times and shifted to EMM-ALU lacking nitrogen, or in EMM-ALU with 0.08% glucose for 90’ before imaging. Sum projections of 5 confocal images acquired over time are shown. Scale bar: 5μm. **B.** Traces of individual Ptc1-GFP fluorescence profiles for n=20 cells grown in EMM-ALU 2% glucose for 24h (left panel) and shifted to 0.08% glucose for 1h before imaging (right panel). Quantified on midplane sum projections of 5 consecutive snapshots. Error bars are standard deviations.

**Table S1:** Cell length at division of indicated genotypes

| Genotypes | Cell length at division | p-values (comparison to WT) | N cells |
| --- | --- | --- | --- |
| wildtype | 13.7 ± 1.0 | - | 300 |
| *ptc1-GFP* | 13.8 ± 1.1 | 0.13 | 300 |
| *ptc1*Δ | 14.2 ± 1.1 | 2.1e<sup>-15</sup> | 300 |
| *ptc2*Δ | 13.2 ± 1.4 | 0.014 | 75 |
| *ptc3*Δ | 13.0 ± 1.0 | 2.6e<sup>-7</sup> | 82 |
| *par1*Δ | 14.9 ± 1.4 | 7.3e<sup>-32</sup> | 300 |
| *par2*Δ | 13.9 ± 1.0 | 0.17 | 83 |
| *ppb1*Δ | 14.7 ± 1.5 | 1.4e<sup>-9</sup> | 85 |
| *ppa1*Δ | 12.7 ± 1.0 | 3.1e<sup>-12</sup> | 141 |
| *ppa3*Δ | 14.0 ± 1.1 | 0.003 | 102 |

